# Clustering gene expression time series data using an infinite Gaussian process mixture model

**DOI:** 10.1101/131151

**Authors:** Ian C. McDowell, Dinesh Manandhar, Christopher M. Vockley, Amy K. Schmid, Timothy E. Reddy, Barbara E. Engelhardt

## Abstract

Transcriptome-wide time series expression profiling is used to characterize the cellular response to environmental perturbations. The first step to analyzing transcriptional response data is often to cluster genes with similar responses. Here, we present a nonparametric model-based method, Dirichlet process Gaussian process mixture model (DPGP), which jointly models cluster number with a Dirichlet process and temporal dependencies with Gaussian processes. We demonstrate the accuracy of DPGP in comparison with state-of-the-art approaches using hundreds of simulated data sets. To further test our method, we apply DPGP to published microarray data from a microbial model organism exposed to stress and to novel RNA-seq data from a human cell line exposed to the glucocorticoid dexamethasone. We validate our clusters by examining local transcription factor binding and histone modifications. Our results demonstrate that jointly modeling cluster number and temporal dependencies can reveal novel regulatory mechanisms. DPGP software is freely available online at https://github.com/PrincetonUniversity/DP_GP_cluster.

## Introduction

The analysis of time series gene expression has enabled insights into development (Kim et al., 2001; Arbeitman et al., 2002; Frank et al., 2015), response to environmental stress (Gasch et al., 2000), cell cycle progression (Cho et al., 1998; Spellman et al., 1998), pathogenic infection (Nau et al., 2002), cancer (Whitfield et al., 2002), circadian rhythm (Panda et al., 2002; Storch et al., 2002), and other biomedically important processes. Gene expression is a tightly regulated spatiotemporal process. Genes with similar expression dynamics have been shown to share biological functions (Eisen et al., 1998). Clustering reduces the complexity of a transcriptional response by grouping genes into a small number of response types. Given a set of clusters, genes are often functionally annotated by assuming *guilt by association* (Walker et al., 1999), sharing sparse functional annotations among genes in the same cluster. Furthermore, regulatory mechanisms characterizing shared response types can be explored using these clusters by, for example, comparing sequence motifs or other features within and across clusters.

Gene clustering methods partition genes into disjoint clusters based on the similarity of expression response. Many clustering methods, such as hierarchical clustering (Eisen et al., 1998), k-means clustering (Tavazoie et al., 1999), and self-organizing maps (Tavazoie et al., 1999), evaluate response similarity using correlation or Euclidean distance. These methods assume that expression levels at adjacent time points are independent and identically distributed, which is statistically invalid for transcriptomic time series data (Ramoni et al., 2002). Some of these methods require a prespecified number of clusters, which may require model selection or post-hoc analyses to determine the most appropriate number.

In model-based clustering approaches, similarity is determined by how well the responses of any two genes fit the same generative model (Yeung et al., 2001a; Ramoni et al., 2002). Model-based methods thus have a clear definition of a cluster (Pan et al., 2002), which is a set of genes that is more likely to be generated from a particular cluster-specific model than other possible models. Mclust, for example, assumes a Gaussian mixture model (GMM) to capture the mean and covariance of expression within a cluster. Mclust selects the optimal number of clusters using the Bayesian information criterion (BIC) (Fraley and Raftery, 2002). However, cluster-specific parameter estimates in Mclust do not take into account uncertainty in cluster number (Medvedovic and Sivaganesan, 2002).

To address the problem of cluster number uncertainty, finite mixture models can be extended to in finite mixture models with a Dirichlet process (DP) prior. This infinite mixture model approach is used in the Gaussian Infinite Mixture Model (GIMM) (Medvedovic et al., 2004; Qin, 2006). Using Markov chain Monte Carlo (MCMC) sampling, GIMM iteratively samples cluster-specific parameters and assigns genes to existing clusters or creates a new cluster based on both the likelihood of the gene expression values with respect to the cluster-specific model and the size of each cluster (Medvedovic et al., 2004). An advantage of nonparametric models is that they allow cluster number and parameter estimation to occur simultaneously when computing the posterior. The DP prior has a “rich get richer” property, meaning that clusters are prioritized for inclusion of a new gene in proportion to cluster size, so bigger clusters are proportionally more likely to grow relative to smaller clusters. This allows for varied cluster sizes as opposed to approaches that favor equivalently sized clusters, such as k-means clustering.

Clustering approaches for time series data that encode dependencies across time have also been proposed. SplineCluster models the time-dependency of gene expression data by fitting non-linear spline basis functions to gene expression profiles, followed by agglomerative Bayesian hierarchical clustering (Heard et al., 2006). The Bayesian Hierarchical Clustering (BHC) algorithm also performs Bayesian agglomerative clustering as an approximation to a DP model, merging clusters until the posterior probability of the merged model no longer exceeds that of the unmerged model (Heller and Ghahramani, 2005; Savage et al., 2009; Cooke et al., 2011). Each cluster in BHC is parameterized by a Gaussian process (GP) with a squared exponential kernel. With this greedy approach, BHC does not capture uncertainty in the clustering.

Recently, models combining DPs and GPs have been developed for time series data analysis. For example, a recent method combines the two to cluster low-dimensional projections of gene expression (Rasmussen et al., 2009). The semiparametric Bayesian latent trajectory model was developed to perform association testing for time series responses, integrating over cluster uncertainty (Dunson and Herring, 2006). Other methods using DPs or approximate DPs to cluster GPs for gene expression data make different modeling decisions (Hensman et al., 2015), different parameter inference methods (Savage et al., 2009), or do not include software that can be easily applied by biologists or bioinformaticians (Rasmussen et al., 2009; Hensman et al., 2015).

Here we develop a statistical model for clustering time series data, the Dirichlet process Gaussian process mixture model (DPGP), and we package this model in user-friendly software. specifically, we combine DPs for incorporating cluster number uncertainty and GPs for modeling time series dependencies. In DPGP, we explore the number of clusters and model the time dependency across gene expression data by assuming that genes within a cluster are generated from a GP with a cluster-specific mean function and covariance kernel. A single clustering can be selected according to one of a number of optimality criteria; alternatively, a gene-by-gene matrix can be generated that reflects the estimated probability that each pair of genes is in the same cluster.

To demonstrate the applicability of DPGP to gene expression response data, we applied our algorithm to simulated, published, and original transcriptomic time series data. We first applied DPGP to hundreds of diverse simulated data sets and show favorable comparisons to other state-of-the-art methods for clustering time series data. DPGP was then applied to a previously published microarray time series data set, recapitulating known gene regulatory relationships (Sharma et al., 2012). To enable biological discovery, RNA-seq data were generated from the human lung epithelial adenocarcinoma cell line A549 from six time points after treatment with dexamethasone (dex) for up to 11 hours. By integrating our DPGP clustering results on these data with a compendium of ChIP-seq data sets from the ENCODE project, we reveal novel mechanistic insights into the genomic response to dex.

## Results

### DPGP compares favorably to state-of-the-art methods on simulated data

We tested whether DPGP recovers true cluster structure from simulated time series data. We applied DPGP to 620 data sets generated using a diverse range of cluster sizes and expression traits (***Supplementary file 1***). We compared our results against those from BHC (Savage et al., 2009), GIMM (Medvedovic et al., 2004), hierarchical clustering by average linkage (Eisen et al., 1998), k-means clustering (Tavazoie et al., 1999), Mclust (Fraley and Raftery, 2002), and SplineCluster (Heard et al., 2006). To compare observed partitions to true partitions, we used *Adjusted Rand Index* (ARI), which measures the similarity between a test clustering and ground truth in terms of cluster agreement for element pairs (Rand, 1971; Hubert and Arabie, 1985). ARI is scaled such that it is 1 when two partitions agree exactly and 0 when two partitions agree no more than is expected by chance (Rand, 1971; Hubert and Arabie, 1985). ARI was recommended in a comparison of metrics (Milligan and Cooper, 1986) and has been used to compare clustering methods in similar contexts (Yeung et al., 2001b; Medvedovic et al., 2004; Dahl, 2006; ***Fritsch et al., 2009***).

Assuming GPs as generating functions, we simulated data sets with varied cluster sizes, length scales, signal variance, and noise variance (***Supplementary file 1***). In this collection of simulations, on the task of reproducing a specific clustering, DPGP generally outperformed GIMM, k-means, and Mclust, but was generally outperformed by BHC and SplineCluster, and performed about as well as hierarchical clustering (Figure 1 and ***Supplementary file 2***). The performance of hierarchical clustering and k-means benefited from prespecification of the true number of clusters—with a median number of 24 clusters across simulations—while other methods were expected to discover the true number of clusters. The more poorly performing methods on these particular data—GIMM, k-means, and Mclust do not model temporal dependency, suggesting that there is substantial value in explicitly modeling the time dependence of observations.

**Figure 1.**
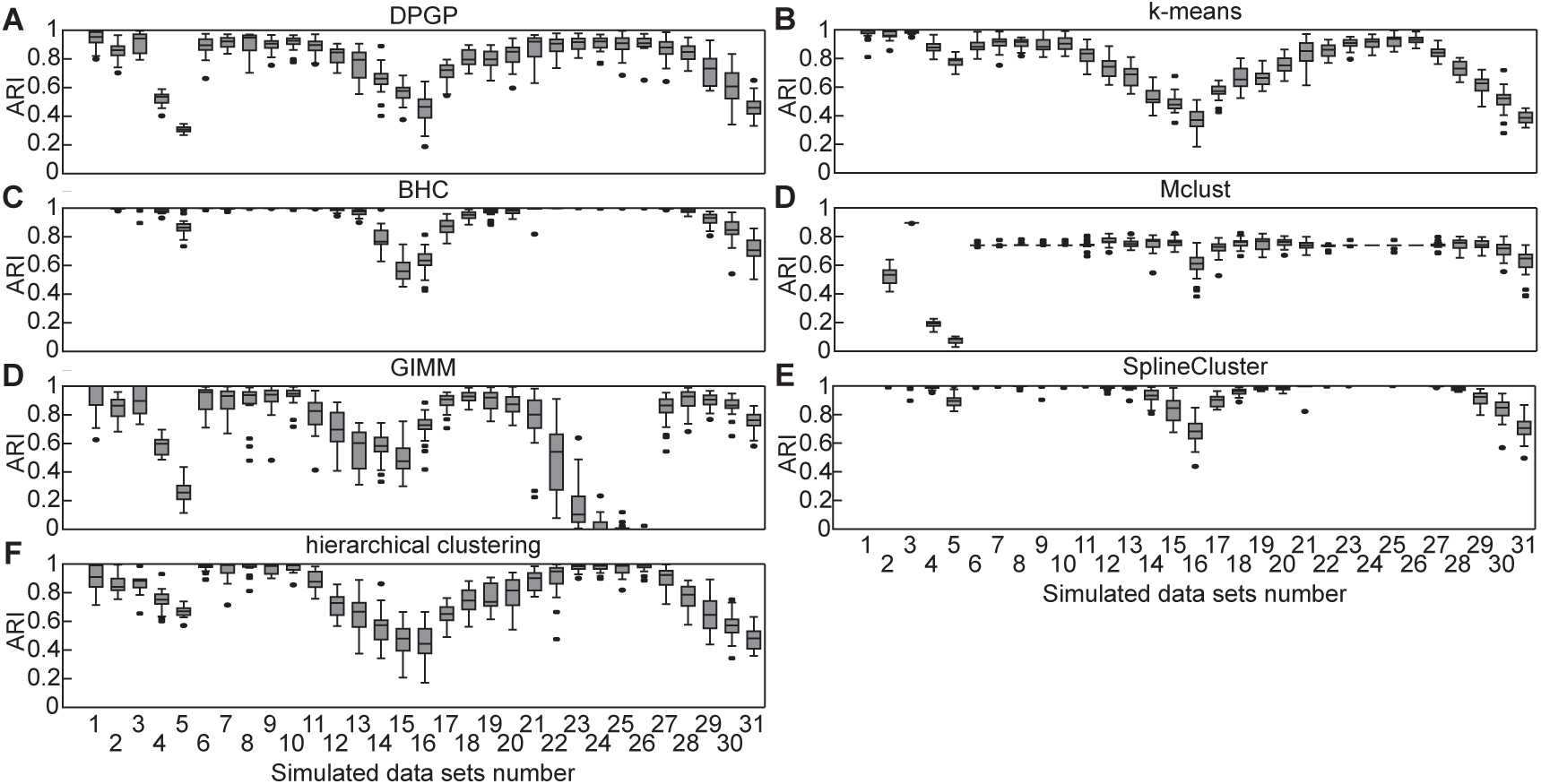
Clustering performance of state-of-the-art algorithms on simulated time series data. Box plots show empirical distribution of clustering performance for each method in terms of Adjusted Rand Index (ARI) across twenty instances of the 31 data set types detailed in ***Supplementary file 1***. Higher values represent better recovery of the simulated clusters. Results shown for (**A**) DPGP, (**B**) k-means, (**C**) BHC, (**D**) Mclust, (**E**) GIMM, (**F**) SplineCluster, and (**G**) hierarchical clustering. **Figure 1–Figure supplement 1.** Time benchmark. Mean runtime of BHC, GIMM, and DPGP across varying numbers of gene expression trajectories generated from GPs parameterized in the same manner as simulated data sets 11, 21, and 27 in ***Supplementary file 1***. Cluster sizes were 2, 4, 8, 16, 32, and 64 for 126 simulated genes in 2–8 different clusters per cluster size. Error bars represent standard deviation in runtime across 20 simulated data sets. Hierarchical clustering, k-means, Mclust, and SplineCluster are not shown because their mean runtimes were under one minute and could not be meaningfully displayed here. **Figure 1–Figure supplement 2.** Proportion of held-out test points within credible intervals of estimated cluster means for DPGP. For all data sets detailed in ***Supplementary file 1***, expression trajectories were clustered while separately holding out each of the four middle time points of eight total time points. Box plot shows proportion of test points that fell within the 95% credible intervals (CIs) of the estimated cluster mean. analyses, similar fractions of genes were found to be directly bound by RosR according to ChIP-chip data from cells exposed to H_2_O_2_ for 0, 10, 20, and 60 minutes (Tonner et al., 2015). When all RosR binding at all four ChIP-chip time points were considered together, 8.9% of DPGP genes changing clusters were bound; 9.5% of DEGs were bound in the previous analysis (Sharma et al., 2012).

DPGP successfully recovered true cluster structure across a variety of generating assumptions except in cases of a large number of clusters each with a small number of genes (data sets 4 and 5) and in cases of small signal variance (data set 16) and high noise variance (data set 31; Figure 1). DPGP was substantially faster than GIMM and BHC, but slower than hierarchical clustering, k-means, Mclust, and SplineCluster (Figure 1–Figure Supplement 1, Wilcoxon two-sided signed-rank, comparing clustering times on 20 data sets with 1, 008 simulated trajectories, DPGP versus each method, *p* ≤ 8.86 × 10^−^5).

An important advantage of DPGP is that—being a probabilistic method uncertainty in clustering and cluster trajectories is modeled explicitly. Some implications of the probabilistic approach are that cluster means and variances can be used to quantify the fit of unseen data, to impute missing data points at arbitrary times, and to integrate over uncertainty in hypothesis testing with the clusters (Dunson and Herring, 2006). Using the same data sets simulated for the algorithm comparison, we clustered expression trajectories while holding out each of the four middle time points of eight total time points. We computed the proportion of held-out test points that fell within the 95% credible intervals (CIs) of the estimated cluster means. For comparison, we also permuted cluster membership across all genes 1, 000 times and recomputed the same proportions. We found that DPGP provided accurate CIs on the simulated gene expression levels (Figure 1–Figure Supplement 2). Across all simulations, at least 90% of test points fell within the estimated 95% CI, except for data set types with large length-scales or high signal variances (both parameters ∈ {1.5, 2, 2.5, 3}). The proportion of test points that fell within the 95% CIs was consistently higher for true clusters than for permuted clusters [Mann-Whitney U-test (MWU), *p* ≤ 2.24 × 10^-6^], except for data with very small length-scales ({0.1,…, 0.5}) in which the proportions were equivalent (MWU, *p* = 0.24). This implies that the simulated sampling rates in these cases were too low for DPGP to capture the temporal patterns in the data.

For the simulations in which DPGP performed worse than BHC or Spline Cluster in recovering the true cluster structure, the clusters inferred from the data provided useful and accurate CIs for unseen data. For example, DPGP performed decreasingly well as the noise variance was increased to 0.4, 0.5, and 0.6. However, the median proportions of test points within the 95% CIs were 93.4%, 92.6%, 91.9%, respectively (Figure 1–**Figure Supplement 2**). This suggests that DPGP provides well calibrated CIs on expression levels over the time course.

We can also use DPGP to evaluate the posterior probability of a specific clustering with respect to the fitted model. Critically, only in 1.6% of all simulated data sets was the posterior probability of the true clustering, given the DPGP model, greater than both the posterior probability of the DPGP MAP partition and than the mean posterior probability across all DPGP samples (Z-test, *p <* 0.05). These results imply that, even in cases where DPGP did not precisely recover the cluster structure, the posterior probability was not strongly peaked around the true partition, meaning that there was substantial uncertainty in the optimal partition.

### Clustering oxidative stress transcriptional responses in a microbial model organism recapitulates known biology

Given the performance of DPGP on simulated data with minimal user input for selection of cluster number, we next sought to assess the performance of DPGP on biological data. As a test case, we applied DPGP to published data from a single-celled model organism with a small genome (*Halobacterium salinarum*, 2.5 Mbp and 2, 400 genes) exposed to oxidative stress induced by addition of H_2_O_2_ (Sharma et al., 2012). This multifactorial experiment tested the effect of deletion of the gene encoding the transcription factor (TF) RosR, which is a global regulator that enables resilience of *H. salinarum* to oxidative stress (Tonner et al., 2015). specifically, transcriptome profiles of a RosR deletion mutant strain (**Δ***rosR*) and control strain were captured with microarrays at 10^-20^ minute intervals following exposure to H_2_O_2_. In the original study, 616 genes were found to be differentially expressed (DEGs) in response to H_2_O_2_, 294 of which were also DEGs in response to RosR mutation. In previous work, the authors clustered those 294 DEGs using k-means clustering with *k* = 8 (minimum genes per cluster = 13, maximum = 86, mean = 49) (Sharma et al., 2012).

We used DPGP on these *H. salinarum* time series data to cluster expression trajectories from the 616 DEGs in each strain independently, which resulted in six clusters per strain (Figure 2). The number of genes in clusters from DPGP varied widely across clusters and strains (minimum 2 genes, maximum 292, mean 102.7) with greater variance in cluster size in trajectories from the mutant strain. To assess how DPGP clustering results compared to previous results using k-means, we focused on how deletion of RosR affected gene expression dynamics. Out of the 616 DEGs, 372 moved from a cluster in the control strain to a cluster with a different dynamic trajectory in **Δ**.*rosR* (e.g., from an up-regulated cluster under H_2_O_2_ in control, such as cluster 5, to a down-regulated cluster in **Δ**.*rosR*, such as cluster 3; Figure 2 and ***Supplementary file 3***). Of these 372 genes, 232 were also detected as differentially expressed in our previous study (Sharma et al., 2012) [significance of overlap, Fisher’s exact test (FET), *p* ≤ 2.2 × 10^-16^]. Comparing these DPGP results to previous

**Figure 2.**
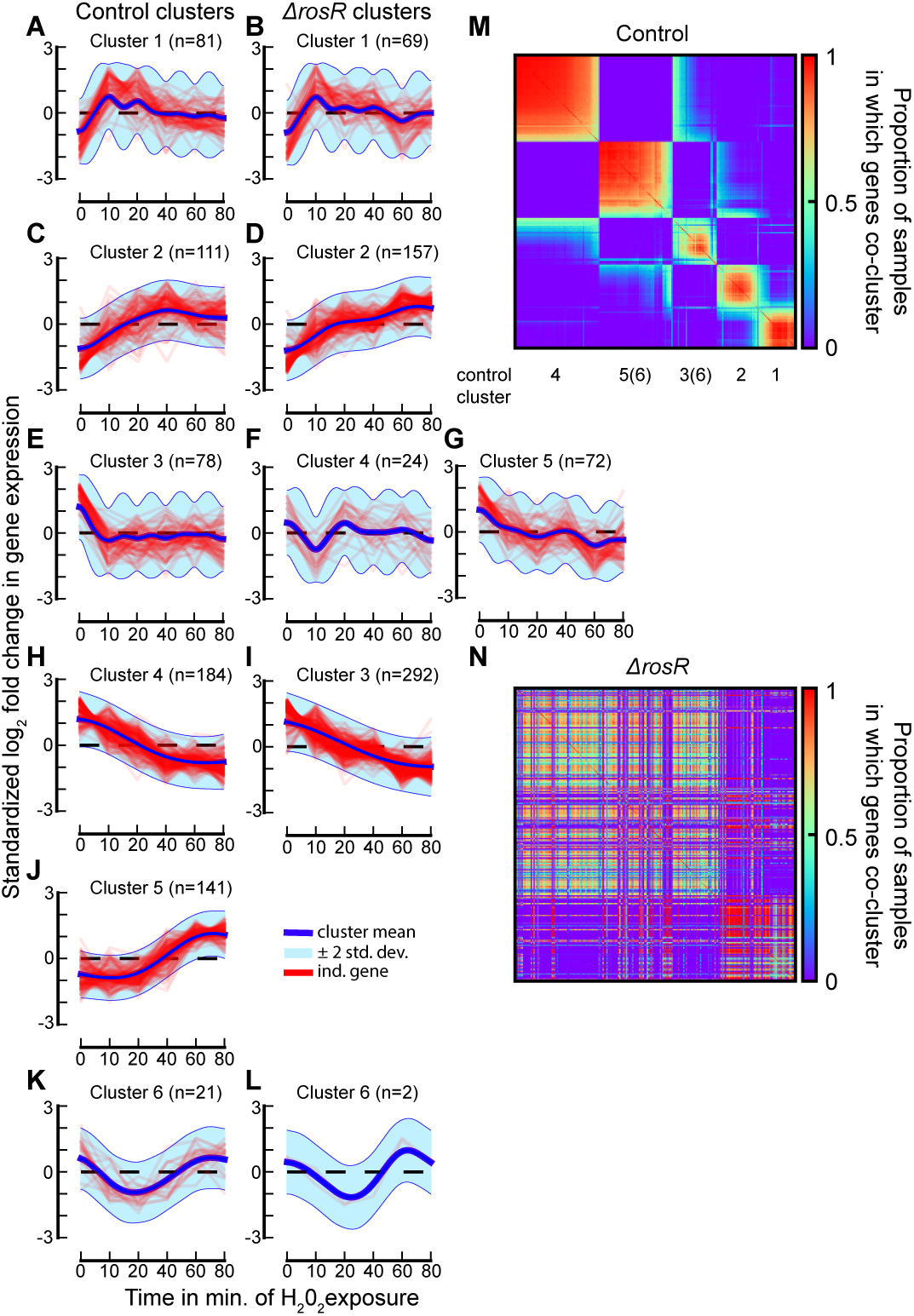
DPGP clusters in *H. salinarum* H_2_O_2_-exposed gene expression trajectories. (**A–L**) For each cluster, standardized log2 fold change in expression from pre-exposure levels is shown for each individual gene as well as the posterior cluster mean ±2 standard deviations. Control strain clusters are on left and **Δ**.*rosR* clusters on right, organized to relate the **Δ**.*rosR* cluster(s) that correspond(s) to each control cluster. (**M**) Heatmap displays the proportion of DPGP samples in which each gene (row/column) clusters with every other gene in the control strain. Rows and columns were clustered by Ward’s linkage. The predominant, clearly visible blocks of elevated co-clustering are labeled with the control cluster numbers to which the genes that compose the majority of the block belong. As indicated, cluster 6 is dispersed across multiple blocks, primarily the blocks for clusters 3 and 5. (**N**) Same as (M), except that values are replaced by the proportions in the **Δ**.*rosR* strain instead of the control strain. Rows and columns ordered as in (M).

Genes most dramatically affected by deletion of *rosR* were those up-regulated after 40 minutes of H_2_O_2_ exposure in the control strain: All 141 genes in control cluster 5 changed cluster membership in the b**Δ**.*rosR* strain (***Figure 2***; FET, *p* ≤ 2.2 × 10^-16^). Of these 141 genes up-regulated in control strains in response to H_2_O_2_, 89 genes (63%) exhibited inverted dynamics, changing to down-regulated in the **Δ**.*rosR* strain. These 89 genes grouped into two clusters in the b.*rosR* strain (**Δ**.*rosR* clusters 3 and 5; Figure 2 and ***Supplementary file 3***). The transcriptional effect of RosR deletion noted here accurately reflects previous observations: 84 of these 89 genes showed differential trajectories in the control versus **Δ**.*rosR* strains previously (Sharma et al., 2012). RosR is required to activate these genes in response to H_2_O_2_ (Sharma et al., 2012). These results suggest that DPGP analysis accurately recapitulates previous knowledge of RosR-mediated gene regulation in response to H_2_O_2_ with substantially reduced user input.

### DPGP reveals mechanisms behind the glucocorticoid transcriptional response in a human cell line

Given the performance of DPGP in recapitulating known results for biological data, we next used DPGP for analysis of novel time series data. specifically, we used DPGP to identify co-regulated sets of genes and candidate regulatory mechanisms in the human glucocorticoid (GC) response. GCs, such as dex, are among the most commonly prescribed drugs for their anti-inflammatory and immunosuppressive effects (Hsiao et al., 2007). GCs function in the cell primarily by affecting gene expression levels. Briefly, GCs diffuse freely into cells where they bind to and activate the glucocorticoid receptor (GR). Once bound to its ligand, the GR translocates into the nucleus where it binds DNA and regulates expression of target genes. The induction of expression from GC exposure has been linked to GR binding (Reddy et al., 2009; Pan et al., 2011). However, while there are a plethora of hypotheses regarding repression and a handful of well-studied cases (De Bosscher and Haegeman, 2009; Santos et al., 2011), it has proved difficult to associate repression of gene expression levels with genomic binding on a genome-wide scale (Reddy et al., 2009; Pan et al., 2011). Further, GC-mediated expression responses are far more diverse than simple induction or repression, motivating a time course study of these complex responses (Balsalobre et al., 2000; Biddie and Hager, 2009; John et al., 2009; Stavreva et al., 2012; Vockley et al., 2016).

To characterize the genome-wide diversity of the transcriptional response to GCs and to reveal candidate mechanisms underlying those responses, we performed RNA-seq in the human lung adenocarcinoma-derived A549 cell line after treatment with the synthetic glucocorticoid (GC) dex for 1, 3, 5, 7, 9, and 11 hours, resulting in six time points. This data set is among the most densely sampled time series of the dex-mediated transcriptional response in a human cell line.

DPGP clustered differentially expressed transcripts into four predominant clusters. We used DPGP to cluster 1, 216 transcripts that were differentially expressed at two consecutive time points (FDR ≤ 0.1). DPGP found 13 clusters with a mean size of 119 transcripts and a standard deviation of 108 transcripts (Figure 3 and Figure 3–**Figure Supplement 1**). In order to analyze the shared mechanisms underlying expression dynamics for genes within a cluster and validate cluster membership, we chose to validate the four largest clusters using a series of complementary analyses and data. These four clusters included 74% of the dex-responsive transcripts. We designated these clusters *up-reg-slow*, *down-reg-slow*, *up-reg-fast*, and *down-reg-fast* (Figure 3) where *fast* clusters had a maximal first-order difference in expression between 1 and 3 hours and *slow* clusters had a maximal first-order difference between 3 and 5 hours. A variety of other clusters were identified with diverse dynamics, revealing the complexity of the GC transcriptional response (Figure 3).

**Figure 3.**
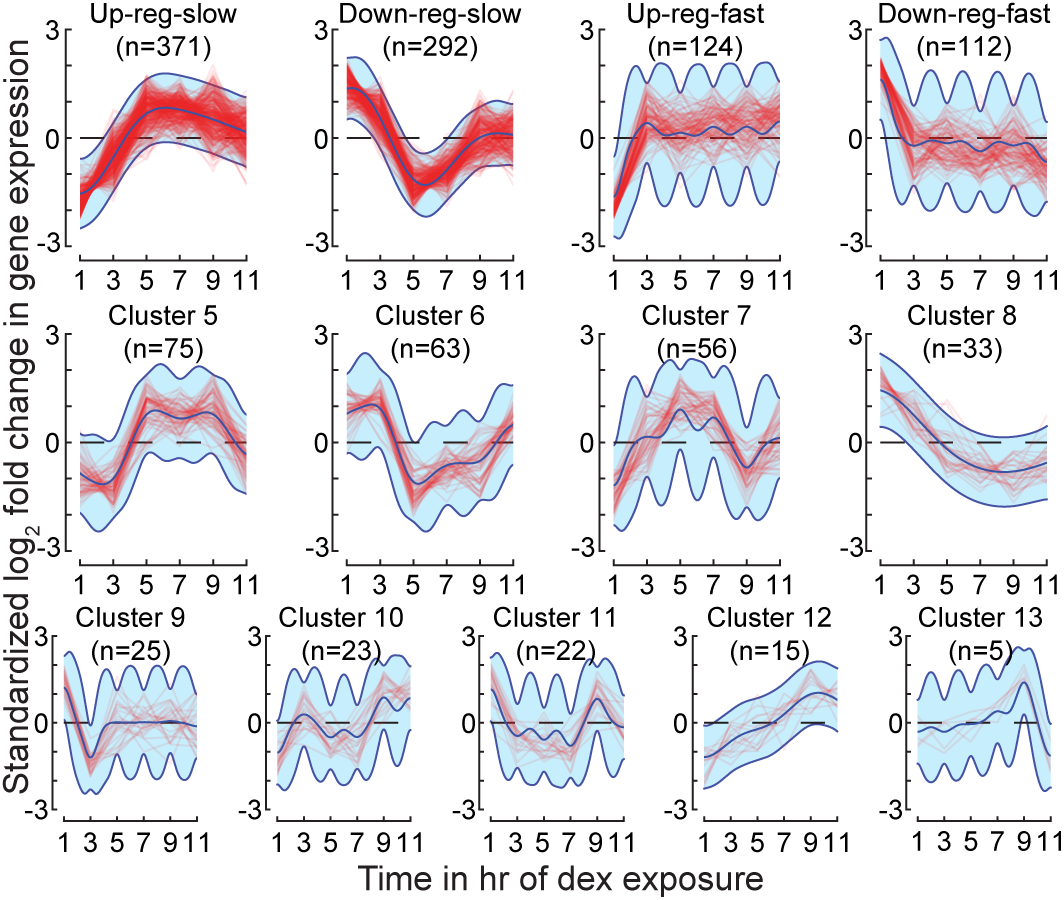
Clustered trajectories of differentially expressed transcripts in A549 cells in response to dex. For each cluster, standardized log2 fold change in expression from pre-dex exposure levels is shown for each transcript, and the posterior cluster mean and ±2 standard deviations according to the cluster-specific GP. **Figure 3–Figure supplement 1.** Rugplot of all cluster sizes for A549 glucocorticoid exposure data clustered using DPGP. Each stick on the x-axis represents a singular data cluster of the 13 total clusters. Note that the two clusters with sizes 22 and 23 are difficult to distinguish by eye.

DPGP dex-responsive expression clusters differ in biological processes.

Genes involved in similar biological processes often respond similarly to stimuli (Eisen et al., 1998). To determine if the DPGP clusters were enriched for genes that contribute to distinct biological processes, we tested each cluster for enrichment of Gene Ontology slim (GO-slim) biological process terms (Ashburner et al., 2000). The *down-reg-slow* cluster was enriched for cell cycle-related terms such as *cell cycle*, *cellular aromatic compound metabolic process*, *heterocycle metabolic process*, *chromosome segregation*, and *cell division*, among other associated terms (see ***Supplementary file 4*** for p-values). This cluster included genes critical to cell cycle progression such as *BRCA2*, *CDK1*, *CDK2*, and others. The down-regulation of these genes is consistent with the antiproliferative effects of GCs (Goya et al., 1993; Rogatsky et al., 1997; King and Cidlowski, 1998). In contrast, the *down-reg-fast* cluster was enriched for terms related to *developmental process* such as *anatomical structure formation involved in morphogenesis* and other terms (***Supplementary file 4***). Genes in the *down-reg-slow* cluster that were annotated as *anatomical structure formation involved in morphogenesis* included homeobox genes like *EREG*, *HNF1B*, *HOXA3*, and *LHX1* as well as growth factors like *TGFA* and *TGFB2*. Our results suggest that GC exposure in A549 cells leads to a rapid down-regulation of growth-related TFs and cytokines and a slower down-regulation of crucial cell cycle regulators.

The *up-reg-slow* and *up-reg-fast* clusters did not differ substantially in functional enrichment, and both were enriched for *signal transduction*. Up-regulated genes annotated as *signal transduction* included multiple MAP kinases, *JAK1*, *STAT3* and others. Whereas the *down-reg-slow* cluster was enriched for genes annotated as *heterocycle metabolic process*, the *up-reg-slow* cluster was depleted (***Supplementary file 3***). Overall, clustering enabled improved insight into GC-mediated transcriptional responses. Our results suggest that a novel functional distinction may exist between rapidly and slowly down-regulated genes.

DPGP clusters differ in TF and histone modification occupancy prior to dex exposure. We validated the four major expression clusters by identifying distinct patterns of epigenomic features that may underlie differences in transcriptional response to GC exposure. In particular,we looked to see whether the co-clustered genes had similar TF binding and chromatin marks before dex exposure. We hypothesized that similar transcriptional responses were driven by similar regimes of TF binding and chromatin marks. To test this, we used all ChIP-seq data generated by the ENCODE project (***Consortium et al., 2012***) that were assayed in the same cell line and treatment conditions (***Supplementary file0 5***). For each data set and each transcript, we counted pre-aligned ChIP-seq reads in three bins of varied distances from the transcription start site (TSS; *<* 1 kb, 1–5 kb, 5–20 kb), based on evidence that suggests that different TFs and histone modifications function at different distances from target genes (Garber et al., 2012). Both TF binding and histone modification occupancy are well correlated (Heintzman et al., 2009; Cheng and Gerstein, 2011). In order to predict cluster membership of each transcript based on a parsimonious set of TFs and histone modifications in control conditions, we used elastic net regression, which tends to include or exclude groups of strongly correlated predictors using a regularized model (Zou and Hastie, 2005). We controlled for differences in basal expression prior to dex exposure by including the baseline transcription level as a covariate in the model.

The features that were most predictive of cluster membership—indicating an association with expression dynamics—were distal H3K36me3, promoter-proximal E2F6, and distal H3K4me1 (Figure 4A, Figure 4–**Supplemental Figure 1**). H3K36me3 marks the activity of transcription, and is deposited across gene bodies, particularly at exons (Krogan et al., 2003; Kolasinska-Zwierz et al., 2009). Its strength as a predictor of cluster membership may represent differences in the methylation of H3K36 between clusters of genes or, alternatively, residual differences in basal expression. E2F6 functions during G1/S cell cycle transition (Bertoli et al., 2013) and its binding was greater in the *down-reg-slow* cluster, which is consistent with the enrichment of genes with cell cycle biological process terms in the same cluster. H3K4me1 correlates strongly with enhancer activity (Heintzman et al., 2009) and the negative coefficient in our model for the down-reg-slow cluster suggests that the contribution of enhancers to expression differs across clusters (Figure 4).

**Figure 4.**
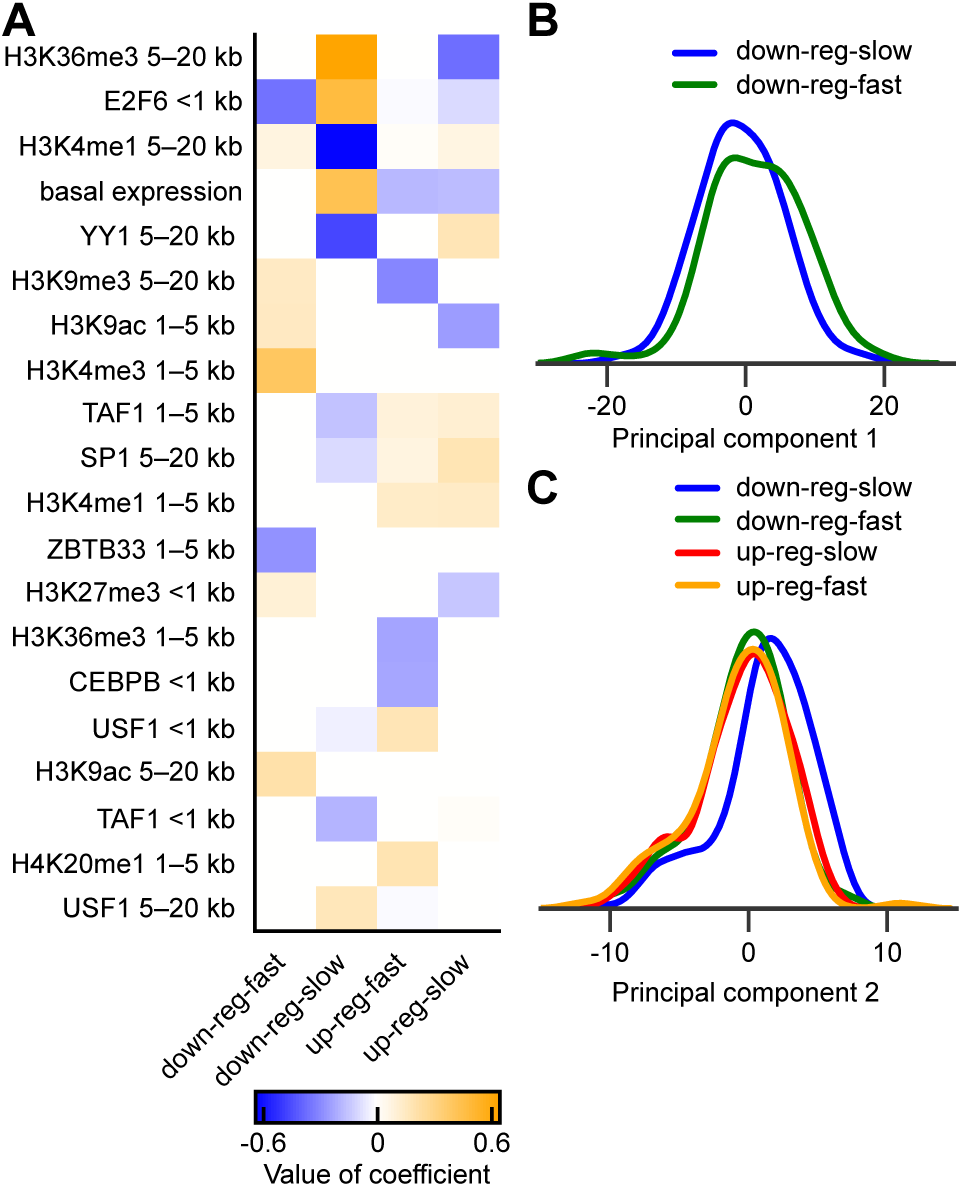
Differences in TF binding and histone modification occupancy in A549 cells in control conditions for the four largest DPGP clusters. (**A**) Heatmap shows the elastic net logistic regression coefficients for the top twenty predictors (sorted by sum of absolute value across clusters) of cluster membership for the four largest clusters. Predictors were log10 library size-normalized binned counts of ChIP-seq TF binding and histone modification occupancy in control conditions. Distance indicated in row names represents the bin of the predictor (e.g., *<* 1 kb means within 1 kb of the TSS). An additional 23 predictors with smaller but non-zero coefficients are shown in ***Figure 4–Supplemental Figure 1***. (**B**) Kernel density histogram smoothed with a Gaussian kernel and Scott’s bandwidth (Scott, 1979) of the TF binding and histone modification occupancy log10 library size-normalized binned count matrix in control conditions transformed by the first principal component (PC1) for the two largest down-regulated DPGP clusters. (**C**) Same as (B), but with matrix transformed by PC2 and with the four largest DPGP clusters. **Figure 4–Figure supplement 1.** Heatmap shows all coefficients (sorted by sum of absolute value across clusters) estimated by elastic net logistic regression of cluster membership for the four largest DPGP clusters as predicted by log10 normalized binned counts of ChIP-seq TF binding and histone modifications in control conditions. Distance indicated in row names reflects the bin of the predictor (e.g. *<* 1 kb = within 1 kb of TSS) **Figure 4–Figure supplement 2.** All non-zero coefficients estimated by elastic net logistic regression of cluster membership for two largest down-regulated DPGP clusters on TF binding and histone modifications in A549 cells in control conditions. Distance indicated in row names reflects the bin of the predictor (e.g., 1 kb = within 1 kb of TSS). **Figure 4–Figure supplement 3.** Scree plot of percentage of variance explained by each principal component in decomposition of epigenomic mapping matrix. The log10 normalized ChIP-seq binned counts around the TSS of genes of TF binding and histone modification occupancy in control conditions was decomposed by PCA. The percentage of variance explained by each of the top ten PCs is shown here. (***Figure 4–Figure Supplement 3***). The 42 ChIP-seq covariates with the highest magnitude loadings on PC1 were restricted to distal, non-promoter TF binding and active histone mark occupancy, implicating enhancer involvement (for the value of all loadings on PC1, see ***Supplementary file 6***). specifically, the features with the two highest magnitude loadings on PC1 were both binned counts of distal p300 binding, a histone acetyltransferase that acetylates H3K27 and is well established as an enhancer mark (Visel et al., 2009; Heintzman et al., 2009).

The two large down-regulated clusters differed substantially in TF binding and histone modifications before exposure to dex (Figure 4A). To confirm, we ran the same regression model after limiting prediction to transcripts in those two clusters. We found that distal H3K4me1 and promoter-proximal E2F6 were highly predictive features, and also four distal histone features that have all been associated with enhancer activity (Figure 4–***Figure Supplement 2***) (He et al., 2010; Rada-Iglesias et al., 2011). This analysis suggests predictive mechanistic distinctions between quickly and slowly down-regulated transcriptional responses to GC exposure. When we performed elastic net regression to identify differential epigenomic features across only the two large up-regulated transcript clusters, on the other hand, no TFs or histone marks were differentially enriched across clusters, meaning that no covariates improved log loss by more than one standard error. This is consistent with our functional enrichment results in which the two up-regulated clusters did not differ substantially in biological process terms.

One drawback of our approach for discriminating between clusters by epigenomic features is that covariates are available for only a handful of such epigenomic features for a specific cell type, and these covariates are often highly correlated (Heintzman et al., 2009; Cheng and Gerstein, 2011). In the context of the elastic net, results should be stable upon repeated inclusion of identical predictors in replicated models (Zou and Hastie, 2005). However, the variables identified as predictive may in truth derive their predictiveness from their similarity to underlying causative TFs or histone modifications. To address the problem of correlated predictors, we used a complementary approach to reveal functional mechanisms distinguishing the four major expression clusters. We projected the correlated features of the standardized control TF and histone modification occupancy data onto a set of linearly uncorrelated covariates using principal components analysis (PCA). We then compared the clusters after transforming each gene’s epigenomic mappings by the two principal axes of variation, which were selected according to the scree plot method (Cattell, 1966) (Figure 4–***Figure Supplement 3***).

The first principal component (PC1) explained 47.9% of the variance in the control ChIP-seq data

We next compared the four largest clusters with respect to their projections onto PC1. We found that the *down-reg-slow* cluster differed substantially from the *down-reg-fast* cluster when transformed by PC1 (MWU, *p* ≤ 2.28 × 10^−3^; Figure 4B), while no other pair wise comparison was signi1cant (MWU, *p >* 0.13). These results suggest that, in aggregate, slowly responding down-regulated transcripts have reduced enhancer activity in control conditions relative to quickly responding down-regulated transcripts.

The second principal component (PC2) explained 11.1% of the variance in the control ChIP-seq data (Figure 4–***Figure Supplement 3***). The 21 ChIP-seq features with the greatest contributions to PC2 captured TF binding and active histone modifications within the promoter (***Supplementary File 5***). By comparing the four largest clusters, we found that the *down-reg-slow* cluster differed from all other clusters with respect to PC2 (MWU, *p* ≤ 9.15 × 10^−7^; Figure 4C), and no other cluster differed from another (MWU, *p >* 0.28). These results illustrate that the slowly responding down-regulated transcripts collectively showed enhanced pre-dex promoter activity compared to the other three largest clusters. Transcriptional response clusters show differences in dynamic TF and histone modification occupancy.

We next validated our four largest dynamic expression clusters by examining the within-cluster similarity in changes in TF binding over time. To do this, we computed the log fold change in normalized ChIP-seq counts for all TFs (CREB1, CTCF, FOXA1, GR, and USF1) assayed through ENCODE with and without 1 hr treatment with 100 nM dex (***Consortium et al., 2012***) (***Supplementary file 5***). We again fit an elastic net logistic regression model, this time to identify the changes in TF binding that were predictive of cluster. The most predictive features of cluster membership were changes in CREB1, FOXA1, and USF1 binding 5–20 kb from the TSS (Figure 5A). CREB1, FOXA1, and USF1 are all known transcriptional activators (Mayr and Montminy, 2001; Corre and Galibert, 2005; Lupien et al., 2008).

**Figure 5.**
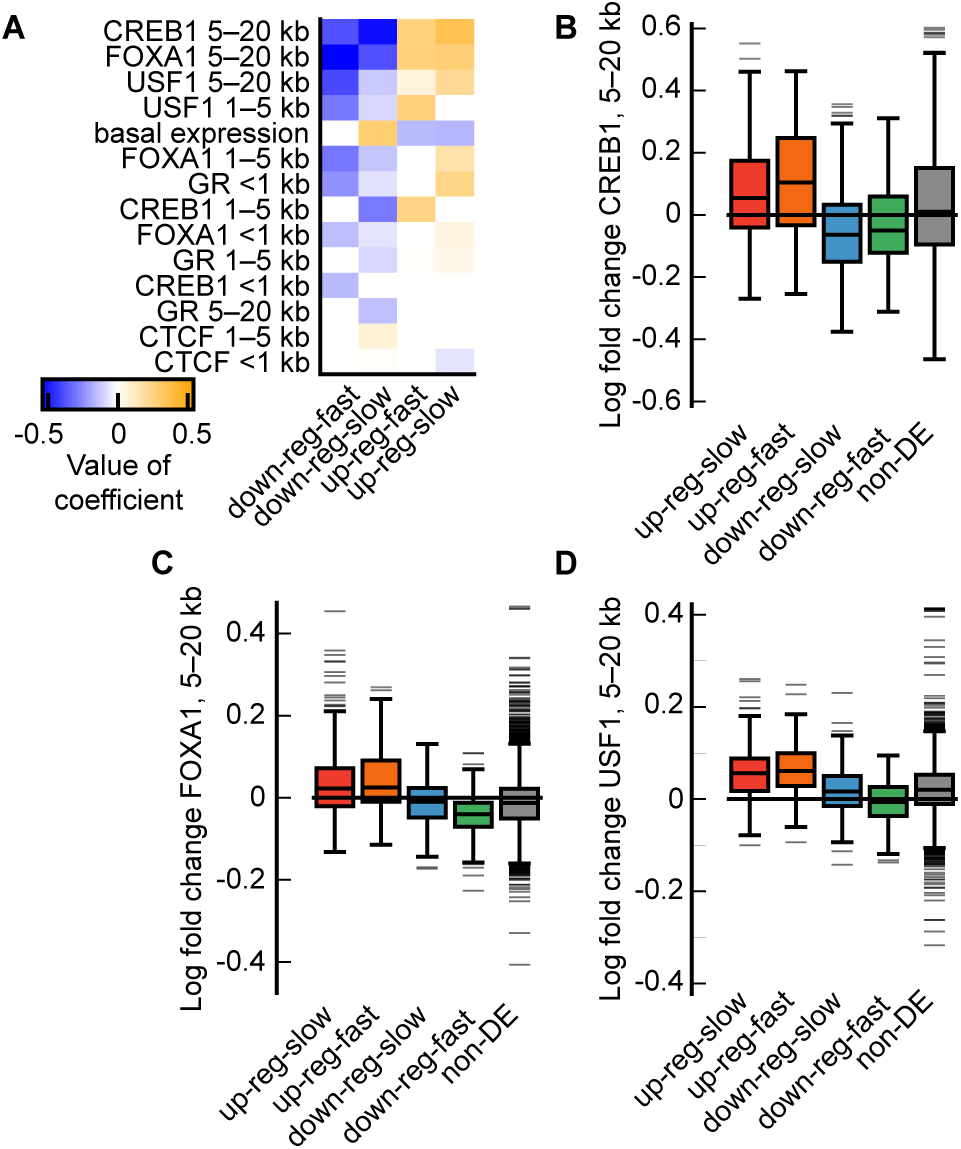
Differences in changes in transcription factor binding in A549 cells in response to glucocorticoid exposure for the four largest DPGP clusters. (**A**) Heatmap shows all coefficients (sorted by sum of absolute value across clusters) for predictors with non-zero coefficients as estimated by elastic net logistic regression of cluster membership for the four largest DPGP clusters. Predictors on y-axis represent log fold-change in normalized binned counts of TF binding from ethanol to dex conditions as assayed by ChIP-seq. Distance indicated in row names reflects the bin of the predictor (e.g. 1 kb = within 1 kb of TSS). (**B**) Cumulative distribution function shows the distances from the TSSs of clustered and non-differentially expressed (non-DE) transcripts to the nearest discrete GR binding peak in dex conditions. (**C**) Boxplots show the logFC in normalized binned counts across clusters and for the group of non-DE transcripts for CREB1, (**D**) FOXA1, and (**E**) USF1.

We examined GR, CREB1, FOXA1, and USF1 binding individually to identify 1ne differences in dynamic TF binding between clusters and compared to stably expressed transcripts. Genes in both up-regulated clusters were closer to the nearest GR binding site. up-regulated clusters had higher median log fold change in binding of the three TFs compared to the two down-regulated clusters (MWU, *p* ≤ 1.5 × 10^-9^, Figure 5B–D). We also noted a number of differences between slowly and quickly responding transcripts. Down-regulated clusters had lower median log fold change in the binding of certain TFs than the group of non-DE transcripts (CREB1 *down-reg-slow* versus non-DE, MWU, *p* ≤ 2.07 × 10^-15^, Figure 5C; CREB1, FOXA1, and USF1 *down-reg-fast* versus non-DE, MWU, *p* = 3.18 × 10^-5^, Figure 5B–D). Additionally, the *down-reg-fast* cluster had lower median log fold change than the *down-reg-slow* cluster in FOXA1 and USF1 binding (MWU, *p* = 8.24 × 10^-6^sa, *p* = 1.29 × 10^-4^, respectively). Overall, increased binding of transcriptional activators was associated with increased expression and with more rapidly increased expression, while decreased binding was associated with decreased expression and more rapidly decreased expression. Our results suggest that differences in TF binding over time may underlie differences in dynamic transcriptional response both in terms of up-regulation versus down-regulation and also in the speed of the transcriptional response.

## Discussion

We developed a Dirichlet process Gaussian process mixture model (DPGP) to cluster measurements of genomic features such as gene expression levels over time. We showed that our method effectively identified disjoint clusters of time series gene expression observations using extensive simulations. DPGP compared favorably to existing methods for clustering time series data and, importantly, includes measures of uncertainty and an accessible, publicly-available software package. We applied DPGP to existing data from a microbial model organism exposed to stress. We found that DPGP accurately recapitulated previous knowledge of TF-mediated gene regulation in response to H_2_O_2_ with minimal user input. We applied DPGP to a novel RNA-seq time series data set detailing the transcriptional response to dex in a human cell line. Our clusters identified four major response types: quickly up-regulated, slowly up-regulated, quickly down-regulated, and slowly down-regulated genes. These response types differed in TF binding and histone modifications before dex treatment and in changes in TF binding following dex treatment.

As with all statistical models, DPGP makes a number of assumptions about observations. In particular, DPGP assumes i) cluster trajectories are stationary; ii) cluster trajectories are exchangeable;each gene belongs to only one cluster; iv) expression levels are sampled at the same time points across all genes; and v) the time point-specific residuals have a Gaussian distribution. Despite these assumptions, our results show that DPGP is robust to certain violations. In the human cell line data, exposure to dex resulted in a non-stationary response (at time point lag 1, all dex-responsive genes had either Augmented Dickey-Fuller *p <* 0.05 or Kwiatkowski–Phillips–Schmidt–Shin, *p >* 0.05), and it has been shown that the residuals may not follow a Gaussian distribution (Schapiro-Wilk test, *p* ≤ 2.2 × 10^−16^), violating assumptions (i) and (v). However, despite these assumption violations, we found that DPGP clustered expression trajectories in a robust and biologically interpretable way. Furthermore, because DPGP does not assume that the gene expression levels are observed at identical intervals within trajectories, DPGP allows study designs with highly irregular sampling across time.

Our DPGP model can be readily extended or interpreted in additional ways. For example, our DPGP returns not only the cluster-specific mean trajectories but also the covariance of that mean, which is useful for downstream analysis by explicitly specifying confidence intervals around interpolated time points. Given the Bayesian framework, DPGP naturally allows for quantification of uncertainty in cluster membership by analysis of the posterior similarity matrix. For example, we could test for association of latent structure with specific genomic regulatory elements after integrating over uncertainty in the cluster assignments (Dunson and Herring, 2006). DPGP can also be applied to time series data from other types of sequencing-based genomics assays such as DNase-seq and ChIP-seq. If we find that the Gaussian assumption is inappropriate for these data types, we may consider using different nonparametric trajectory distributions to model the response trajectories, such as a Student-t process (Shah et al., 2014).

When DPGP was applied to RNA-seq data from A549 cells exposed to GCs, the clustering results enabled several important biological observations. Two down-regulated response types were distinguished from one another based on histone marks and TF binding prior to GC exposure. The rapidly down-regulated cluster included homeobox TFs and growth factor genes and was enriched for enhancer regulatory activity, while slowly down-regulated cluster included critical cell cycle genes and was enriched for promoter regulatory activity. More study is need to resolve how GCs differentially regulate these functionally distinct classes of genes. GR tends to bind distally from promoters (Reddy et al., 2009) so that rapid down-regulation may be a direct effect of GR binding, while slower down-regulation may be secondary effect. We also found that down-regulated genes lost binding of transcriptional activators in distal regions while up-regulated genes gained binding. This result links genomic binding to GC-mediated repression on a genome-wide scale. With increasing availability of high-throughput sequencing time series data, we anticipate that DPGP be a powerful tool for defining cellular response types.

## Materials and methods

### Dirichlet Process Gaussian Process (DPGP) mixture model

We developed a Bayesian nonparametric model for time series trajectories *Y* ER *P* ×*T*, where *P* is the number of genes and *T* the number of time points per sample, assuming observations at the same time points across samples and no missing data. In particular, let *yj* be the vector of gene expression values for gene *j* ∈ {1,…, *P*} for all assayed time points *t* ∈ {1,…, *T*}.Then, we define the generative DP mixture model as follows:

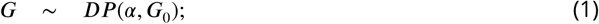

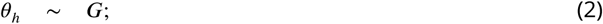

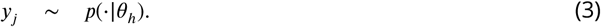

Here, *DP* represents a draw *G* from a DP with *base distribution G*0. *G*, then, is the distribution from which the latent variables **θ***h* are generated for cluster *h*, with **а** *>* 0 representing the *concentration parameter*, with larger values of *α* encouraging more and smaller clusters. We specify the observation distribution *y*_*j*_ ˜ *p* (·| θ_*h*_) with a Gaussian process. With the DP mixture model, we are able to cluster the trajectory of each gene over time without specifying the number of clusters *a priori*.

We can integrate out *G* in the DP to find the conditional distribution of one cluster-specific random variable **θ***h* conditioned on all other variables **θ**¬*h*, which represent the cluster-specific parameter values of the observation distribution (here, a GP); using exchangeability, for all clusters *h* ∈ {1,…, *H*} we have

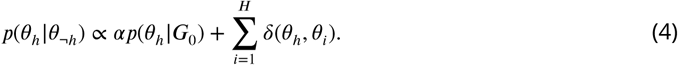

A prior could be placed on **а**, and the posterior for **а** could be estimated conditioned on the observations. Here we favor simplicity and speed, and we set **а** to one. This choice has been used in gene expression clustering (Medvedovic and Sivaganesan, 2002) and other applications (Kim et al., 2006; Vlachos et al., 2008) and favors a relatively small number of clusters, where the expected number of clusters scales as **а** log *P*.

### Gaussian process prior distribution

Our base distribution for the DP mixture model captures the distribution of each parameter of the cluster-specific GP. A GP is a distribution on arbitrary functions mapping points in the input space *xt*—here, time—to a response *yj*—here, gene expression levels of gene *j* across time *t* ∈ {1,…, *T*}. The within-cluster parameters for the distribution of trajectories for cluster *h*, or θ_*h*_ = {*μ*_*h*_, *ℓ*_*h*_ τ_*h*_ 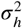}, can be written as follows:

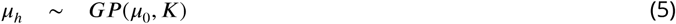

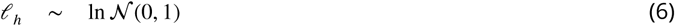

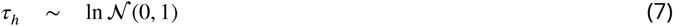

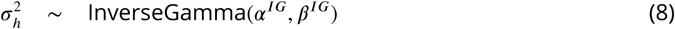

where **а** ^*IG*^ captures shape and **β** ^*IG*^ represents rate (inverse of scale). The above hyperparameters may be changed by the user of the DPGP software. By default, **а** ^*IG*^ is set to 12 and **β** ^*IG*^ is set to 2, as these were determined to work well in practice for our applications. For data with greater variability, such as microarray data, the shape parameter can be decreased to allow for greater noise variance within a cluster. The base distributions of the cluster-specific parameters, which we estimate directly from the data, were chosen to be the natural prior distributions.

The positive definite Gram matrix *Kh* quantifies similarity between every pair of time points *x, x*i in the absence of local noise using Mercer kernel function *K*_*h,t,t‵*_ = *k*_*h*_(*x*_*t*_,*x*_*t‵*_) We used the squared exponential covariance function (dropping the gene index *j*):

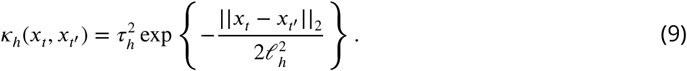

The hyper parameter **ℒ***h*, known as the *characteristic length scale*, corresponds to the distance in input space between two data points smaller than which the points have correlated outputs. The hyperparamete 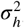 or *signal variance*, corresponds to the variance in gene expression trajectories over time. The model could be easily adapted to different choices of kernel functions depending on the smoothness of the trajectories used in the analysis, such as the Matérn kernel (***Abramowitz et al., 1966***), a periodic kernel (Schölkopf and Smola, 2002), or a non-stationary kernel (Rasmussen and Williams, 2006).

Including local (i.e., time point-specific) noise, *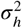* (***Equation 8***), the covariance between time points for trajectory *yj* becomes 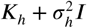. Thus,

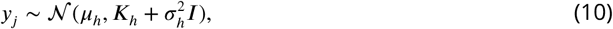

where the noise variance, 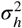, is unique to each cluster *h*. This specifies the probability distribution of each observation *yj* in Equation (3) according to a cluster-specific GP.

### Markov chain Monte Carlo (MCMC) to estimate posterior distribution of DPGP

Given this DPGP model formulation, we now develop methods to estimate the posterior distribution of the model parameters. We use MCMC methods, which have been used previously in time series gene expression analysis (Medvedovic and Sivaganesan, 2002; Qin, 2006). MCMC allows the inference of cluster number and parameter estimation to proceed simultaneously. MCMC produces an estimate of the full posterior distribution of the parameters, allowing us to quantify uncertainty in their estimates. For MCMC, we calculate the probability of the trajectory for gene *j* belonging to cluster *h* according to the DP prior with the likelihood that gene *j* belongs to class *h* according to the cluster-specific GP distribution. We implemented Neal’s Gibbs Sampling “Algorithm 8” to estimate the posterior distribution of the trajectory class assignments (Neal, 2000). More precisely, let *ej* be a categorical latent variable specifying what cluster gene *j* is assigned to, and let *e*¬*j* represent the class assignment vector for all trajectories except for gene *j*. Let *c*_*j*_ represent model parameters where each Ψ_*h*_ = {ℓ_*h*_, τ_*h*_ σ_*h*_} includes parameters specific to cluster *h*.Using Bayes rule, we compute the distribution of each *ej* conditioned on the data and all other cluster assignments:

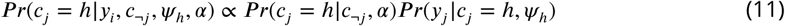

where the first term on the right-hand side represents the probability of assigning the trajectory to cluster *h* and the second term represents the likelihood that the trajectory *yj* was generated from the GP distribution for the *h*th cluster.

According to our model specification, the probability *Pr(c*_*j*_ = *h*|*c*_−*j*_, *α* in Equation (11) is equivalent to the Chinese restaurant process in which:

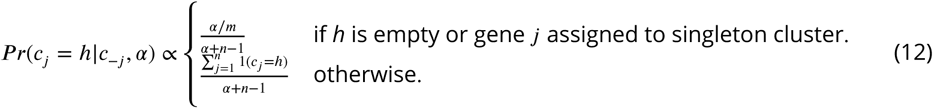

In the above, *m* is the number of empty clusters available in each iteration. Similarly, the likelihood *Pr(y*_*j*_|*c*_*j*_ = *h* ψ_*h*_, *α* in Equation (11) is calculated using our cluster-specific GPs:

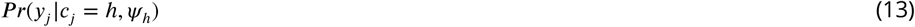

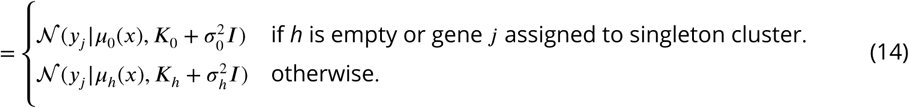

We draw *µ*_0_(*x*) as a sample from the prior covariance matrix, and we put prior distributions on parameters 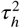, *ℓ*_*h*_ and 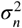 (Equation 6) and estimate their posterior distributions explicitly.

In practice, the first 48% of the prespecified maximum number of MCMC iterations is split into two equally sized burn-in phases. At initialization, each gene is assigned to its own cluster, which is parameterized by its mean trajectory and an SE kernel with unit signal variance and unit length-scale (after the mean time interval between sampling points has been scaled to one unit so that the above length scale hyper prior remains reasonable [***Equation 6***]). The local variance is initialized as the mode of the prior local variance distribution. During the first burn-in phase, a cluster is chosen for each gene at each iteration where the likelihood depends on the fit to a multivariate normal parameterized by the cluster’s mean function and the covariance kernel with initial parameters defined above.

Before each iteration, *m* empty clusters (by default, 4) are re-generated, each of which has a mean function drawn from the prior mean function of 0 with variance equivalent to the noise variance described above. These empty clusters are also assigned the initial covariance kernel parameters described above.

After the second burn-in phase, we update the model parameters for each cluster (at every *sth* iteration to increase speed). specifically, we compute the posterior probabilities of the kernel hyper-parameters. To simplify calculations, we maximize the marginal likelihood, which summarizes model fit while integrating over the parameter priors, known as type II maximum likelihood (Rasmussen and Williams, 2006). specifically:

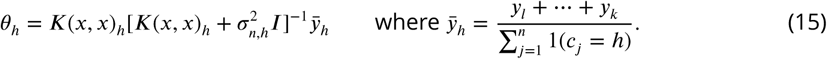

We do this using the fast quasi-Newton limited-memory Broyden-Fletcher-Goldfarb-Shanno (L-BFGS) method implemented in SciPy (Jones et al., 2015). After the second burn-in phase, the cluster assignment vector *c* is sampled at every *sth* iteration to thin the Markov chain, where *s* = 3 by default.

### Selecting the clusters

Our MCMC approach produces a sequence of states drawn from a Gibbs sampler, where each state captures a partition of genes into disjoint clusters. In DPGP, we allow several choices for summarizing results from the Markov chain. Here, we take the maximum a posteriori (MAP) clustering, or the partition that produces the maximum value of the posterior probability. We also summarized the information contained in the Gibbs samples into a *posterior similarity matrix* (PSM) of dimension *P* × *P*, for all genes *P*, where *S*[*j, j*‵] = the proportion of Gibbs samples for which a pair of genes *j, j*‵ are in the same partition, 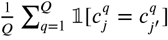 for *Q* iterations of a Gibbs sample and 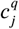 representing the cluster assignment of gene *j* in iteration *q*. This PSM avoids the problem of label switching by being agnostic to the cluster labels when checking for equality.

### Generating the simulated data

In order to test our algorithm across a wide variety of possible data sets, we formulated over twenty generative models with different numbers of clusters (10-100) and with different generative covariance parameters (signal variance 0.5-3, noise variance 0.01-1, and length-scale 0.5-3). We varied cluster number and covariance parameters both across models and within models. For each model, we generated 20 data sets to ensure that results were robust to sampling. In total, we simulated 620 data sets for testing. To generate each data set, we specified the total number of clusters and the number of genes in each cluster. For each cluster, we drew the cluster’s mean expression from a multivariate normal with mean zero and covariance equivalent to a noisy squared-exponential kernel with prespecified hyperparameter settings, then drew a number of samples (gene trajectories) from a multivariate normal with this expression trajectory as mean and the posterior covariance kernel as covariance.

We compared results of DPGP applied to these simulated data sets against results from 1ve state-of-the-art methods, including two popular correlation-based methods, and three model-based methods that use a finite GMM, an infinite GMM, and spline functions, respectively.

- BHC (v.1.22.0) (Savage et al., 2009);
- GIMM (v.3.8) (Medvedovic et al., 2004);
- hierarchical clustering by average linkage (Eisen et al., 1998) (AgglomerativeClustering implemented in SciKitLearn (Pedregosa et al., 2011));
- k-means clustering (Tavazoie et al., 1999) (KMeans, implemented in SciKitLearn (Pedregosa et al., 2011));
- Mclust (v.4.4) (Fraley and Raftery, 2002);
- SplineCluster (v. Oct. 2010) (Heard et al., 2006).

Hierarchical clustering and k-means clustering were parameterized to return the true number of clusters. All of the above algorithms, including our own, were run with default arguments. The only exception was GIMM, which was run by specifying “complete linkage”, so that the number of clusters could be chosen automatically by cutting the returned hierarchical tree at distance 1.0, as in “Auto” IMM clustering (Medvedovic et al., 2004).

We evaluated the accuracy of each approach using ARI. To compute ARI, let *a* equal the number of pairs of co-clustered elements that are in the same true class, *b* the number of pairs of elements in different clusters that are in different true classes, and *N* the total number of elements clustered:

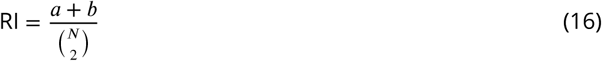

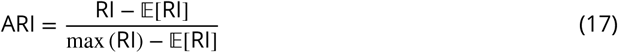

For a derivation of the expectation of RI above, see (Hubert and Arabie, 1985).

### Transcriptional response in *H. salinarum* control strain versus Δ.*rosR* transcription factor knockout in response to H_2_O_2_

Gene expression microarray data from our previous study (Sharma et al., 2012) (GEO accession GSE33980) was clustered using DPGP. In the experiment, *H. salinarum* control and **Δ**.*rosR* TF deletion strains were grown under standard conditions (rich medium, 37C, 225 r.p.m. shaking) until mid-logarithmic phase. Expression levels of all 2,400 genes in the *H. salinarum* genome (Ng et al., 2000) were measured in biological duplicate, each with 12 technical replicate measurements, immediately prior to addition of 25 mM H_2_O_2_ and at 10, 20, 40, 60, and 80 min after addition. The mean of expression across replicates was standardized to mean 0 and variance 1 across all time points and strains. Standardized expression trajectories of 616 non-redundant genes previously identified as differentially expressed in response to H_2_O_2_ (Sharma et al., 2012) were then clustered using DPGP with default arguments, except that the 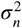 hyperprior parameters were set to **α** IG = 6 and **β** ^*IG*^ = 2 to allow modeling of increased noise in microarray data relative to RNA-seq. Gene trajectories for each of the control and **Δ**.*rosR* strains were clustered in independent DPGP modeling runs. Resultant clusters were analyzed to determine how each gene changed cluster membership in response to the **Δ**.*rosR* mutation. We computed the Pearson correlation coefficient in mean trajectory between all control clusters and all b.*rosR* clusters. Clusters with the highest coefficients across conditions were considered equivalent across strains (e.g., control cluster 1 versus **Δ**.*rosR* cluster 1, *ρ* = 0.886 in Figure 2). significance of overrepresentation in cluster switching (e.g., from control cluster 1 to **Δ**.*rosR* cluster 2) was tested using FET. To determine the degree of correspondence between DPGP results and previous clustering results with the same data, we took the intersection of the list of 372 genes that changed cluster membership according to DPGP with genes in each of eight clusters previously detected using k-means (Sharma et al., 2012). significance of overlap between gene lists was calculated using FET.

### GC transcriptional response in a human cell line

A549 cells were cultured and exposed to the GC dex or a paired vehicle ethanol (EtOH) control as in previous work (Reddy et al., 2009) with triplicates for each treatment and time point. Total RNA was harvested using the Qiagen RNeasy miniprep kit, including on column DNase steps, according to the manufacturer’s protocol. RNA quality was evaluated using the Agilent Tape station and all samples had a RNA integrity number *>* 9. Stranded Poly-A+ RNA-seq libraries were generated on an Apollo 324 liquid handling platform using the Wafergen poly-A RNA purification and RNA-seq kits according to manufacturer instructions. The resulting libraries were then pooled in equimolar ratios and sequenced on two lanes 50 bp paired end lanes on an Illumina HiSeq 2000.

RNA-seq reads were mapped to GENCODE (v.19) transcripts using Bowtie (v.0.12.9) (Langmead et al., 2009) and quantified using samtools idxstats utility (v.1.3.1) (Li et al., 2009). Differentially expressed (DE) transcripts were identified in each time point separately using DESeq2 (v.1.6.3) (Love et al., 2014) with default arguments and FDR ≤ 10%. We clustered only one transcript per gene, in particular, the transcript with the greatest differential expression over the time course among all transcripts for a given gene model, using Fisher’s method of combined p-values across time points. Further, we only clustered transcripts that were differentially expressed for at least two consecutive time points, similar to the approach of previous studies (Nau et al., 2002; Shapira et al., 2009). We standardized all gene expression trajectories to have zero mean and unit standard deviation across time points. We clustered transcripts with DPGP with default arguments.

To query the function of our gene expression clusters, we annotated all transcripts tested for differential expression with their associated biological process Gene Ontology slim (GO-slim) (***Ashburner et al., 2000***) terms and performed functional enrichment analysis using FET with FDR correction (Benjamini and Hochberg, 1995) as implemented in goatools (Tang et al., 2016). We considered results significant with FDR ≤ 5%.

We performed principal components analysis (as implemented in SciKitLearn (Pedregosa et al., 2011)) on the standardized log10 library size-normalized binned counts of TF binding and histone modifications in control conditions only for the observations that corresponded to transcripts in the four largest DPGP clusters.

## Acknowledgments

We thank our Alejandro Barrera who provided insight on packaging the DPGP software. We thank our colleagues at Duke and Princeton Universities for insightful conversations about the research.

## Additional information

### Competing interests

The authors declare that no competing interests exist.

### Funding

BEE was funded by NIH R00 HG006265, NIH R01 MH101822, NIH U01 HG007900, and a Sloan Faculty Fellowship. CMV, ICM, and TER were funded by NIH U01 HG007900. CMV was also funded by NIH F31 HL129743. DM was funded by CBB TG NIH 5T32GM071340. AKS was funded by NSF MCB 1417750. The funders had no role in study design, data collection and analysis, decision to publish, or preparation of the manuscript.

### Author contributions

TER, AKS, and CMV conceived and designed the biological experiments. BEE, ICM, and DM conceived and designed the algorithm and the computational experiments. ICM performed the analysis. ICM, BEE, and AKS analyzed the results. ICM, BEE, TER, and AKS wrote the paper. TER and BEE supervised and funded the research.

## Additional files

### Supplementary files

- Supplementary file 1. Simulated data sets used for algorithm comparisons.
- Supplementary file 2. P-values for algorithm comparisons on simulated data.
- Supplementary file 3. Frequency of switches observed in DPGP clusterings from wild type (**Δ**.*ura*3) to mutant (**Δ**.*rosR*) in H_2_O_2_ exposure in *H. salinarum*.
- Supplementary file 4. Functional enrichment results for four largest DPGP expression clusters in A549 cells in response to the glucocorticoid dexamethasone.
- Supplementary file 5. ENCODE ChIP-seq datasets used in the analysis of GC-responsive clusters.
- Supplementary file 6. Principal components analysis loadings by feature for PC1 and PC2 for ChIP-seq TF binding and histone modifications in A549 cells in control conditions.

---

**Figure 1–Figure supplement 1.**
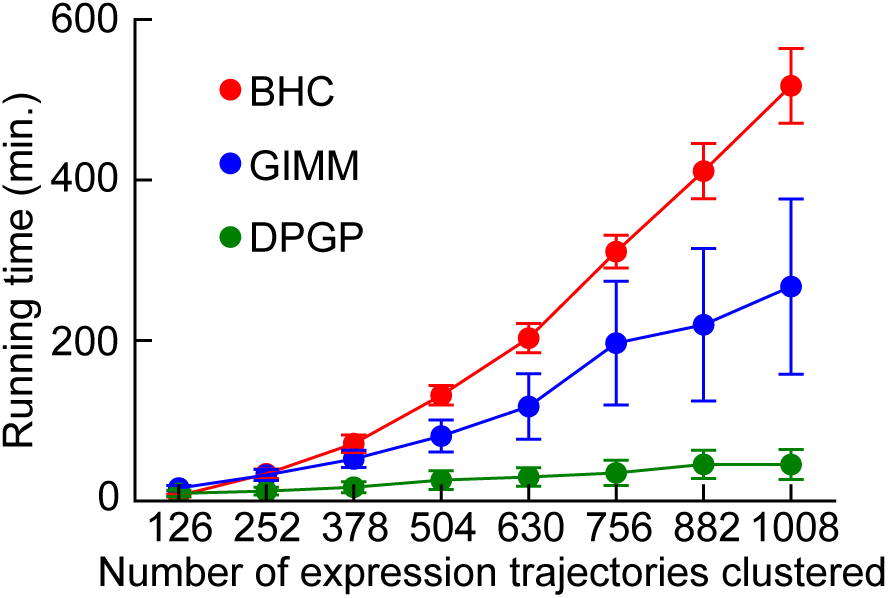
Time benchmark. Mean runtime of BHC, GIMM, and DPGP across varying numbers of gene expression trajectories generated from GPs parameterized in the same manner as simulated data sets 11, 21, and 27 in ***Supplementary file 1***. Cluster sizes were 2, 4, 8, 16, 32, and 64 for 126 simulated genes in 2–8 different clusters per cluster size. Error bars represent standard deviation in runtime across 20 simulated data sets. Hierarchical clustering, k-means, Mclust, and SplineCluster are not shown because their mean runtimes were under one minute and could not be meaningfully displayed here.

**Figure 1–Figure supplement 2.**
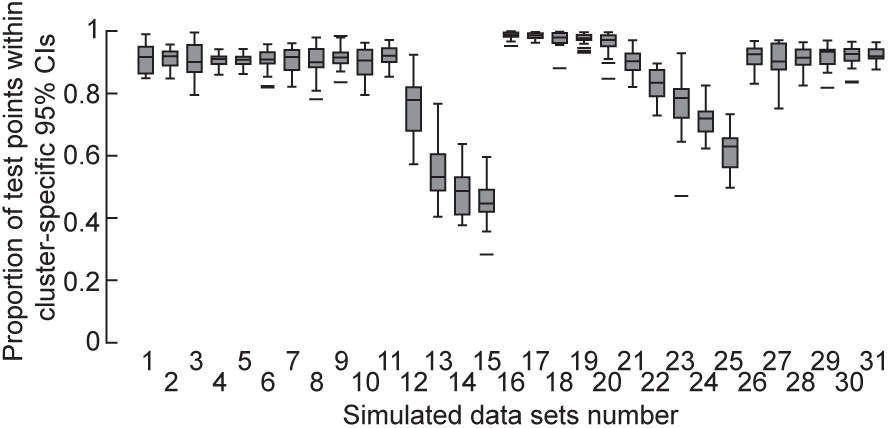
Proportion of held-out test points within credible intervals of estimated cluster means for DPGP. For all data sets detailed in ***Supplementary file 1***, expression trajectories were clustered while separately holding out each of the four middle time points of eight total time points. Box plot shows proportion of test points that fell within the 95% credible intervals (CIs) of the estimated cluster mean.

**Figure 3–Figure supplement 1.**
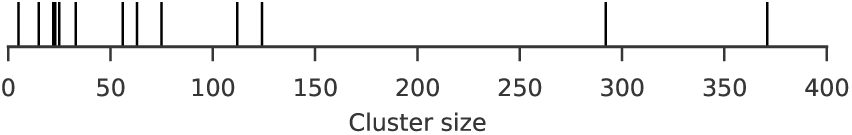
Rugplot of all cluster sizes for A549 glucocorticoid exposure data clustered using DPGP. Each stick on the x-axis represents a singular data cluster of the 13 total clusters. Note that the two clusters with sizes 22 and 23 are difficult to distinguish by eye.

**Figure 4–Figure supplement 1.**
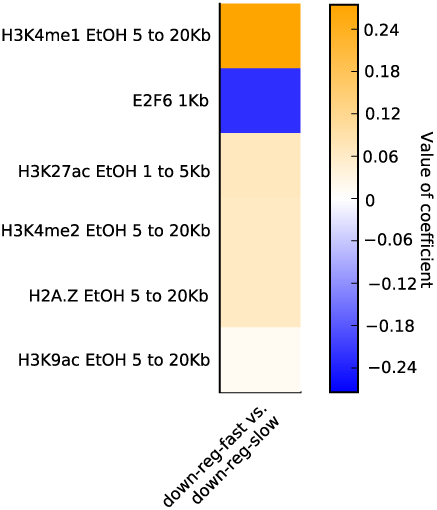
Heatmap shows all coefficients (sorted by sum of absolute value across clusters) estimated by elastic net logistic regression of cluster membership for the four largest DPGP clusters as predicted by log10 normalized binned counts of ChIP-seq TF binding and histone modifications in control conditions. Distance indicated in row names reflects the bin of the predictor (e.g. *<* 1 kb = within 1 kb of TSS)

**Figure 4–Figure supplement 2.**
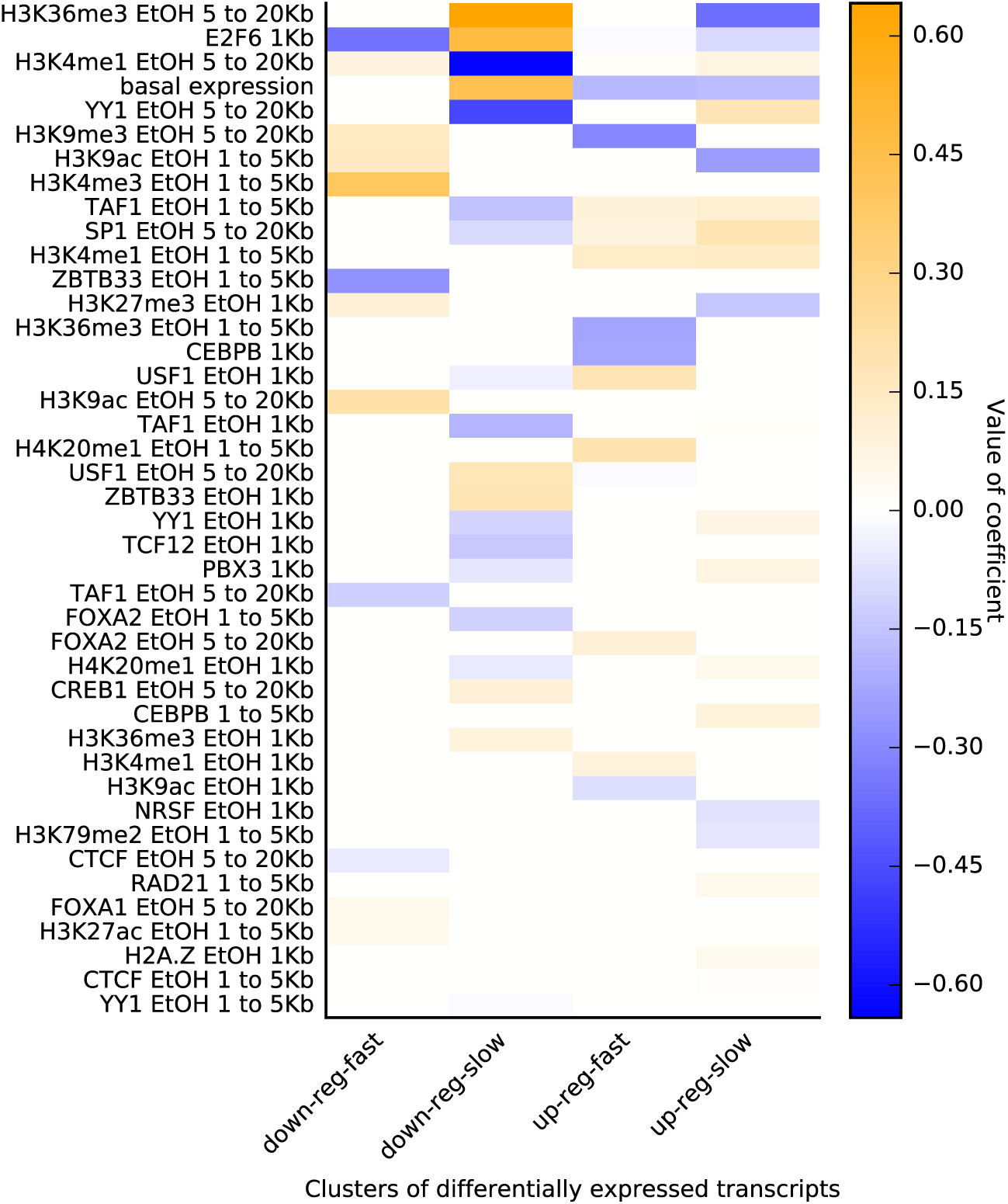
All non-zero coefficients estimated by elastic net logistic regression of cluster membership for two largest down-regulated DPGP clusters on TF binding and histone modifications in A549 cells in control conditions. Distance indicated in row names reflects the bin of the predictor (e.g., 1 kb = within 1 kb of TSS).

**Figure 4–Figure supplement 3.**
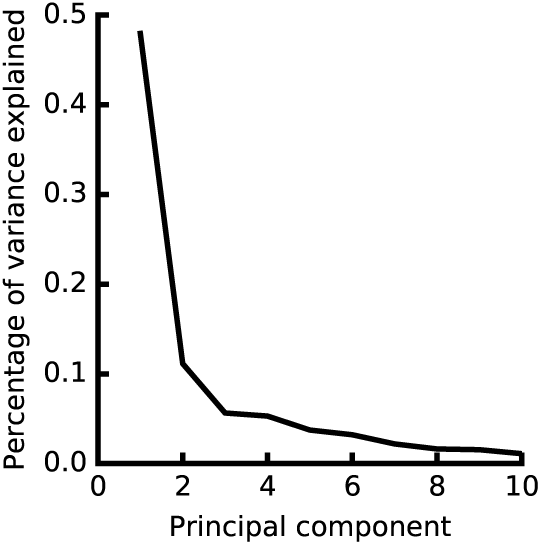
Scree plot of percentage of variance explained by each principal component in decomposition of epigenomic mapping matrix. The log10 normalized ChIP-seq binned counts around the TSS of genes of TF binding and histone modification occupancy in control conditions was decomposed by PCA. The percentage of variance explained by each of the top ten PCs is shown here.

